# Cognitive tasks, anatomical MRI, and functional MRI data evaluating the construct of self-regulation

**DOI:** 10.1101/2023.09.27.559869

**Authors:** Patrick G. Bissett, Ian W. Eisenberg, Sunjae Shim, Jaime Ali H. Rios, Henry M. Jones, Mckenzie P. Hagen, A. Zeynep Enkavi, Jamie K. Li, Jeanette A. Mumford, David P. MacKinnon, Lisa A. Marsch, Russell A. Poldrack

**Affiliations:** Department of Psychology, Stanford University; Credo AI; Department of Psychology, University of Chicago; Department of Psychology, University of Washington; Division of the Humanities and Social Sciences, California Institute of Technology; Department of Psychology, Arizona State University; Center for Technology and Behavioral Health, Geisel School of Medicine, Dartmouth College

## Abstract

We describe the following shared data from N=103 healthy adults who completed a broad set cognitive tasks, surveys, and neuroimaging measurements to examine the construct of self-regulation. The neuroimaging acquisition involved task-based fMRI, resting fMRI, and structural MRI. Each subject completed the following ten tasks in the scanner across two 90- minute scanning sessions: attention network test (ANT), cued task switching, Columbia card task, dot pattern expectancy (DPX), delay discounting, simple and motor selective stop signal, Stroop, a towers task, and a set of survey questions. Subjects also completed resting state scans. The dataset is shared openly through the OpenNeuro project, and the dataset is formatted according to the Brain Imaging Data Structure (BIDS) standard.

## Background & Summary

We report a dataset acquired as part of an effort to understand the construct of *self regulation*, which refers to the processes or abilities that are used to serve long-term goals. Self regulation has been shown to relate to a variety of real-world outcomes, including economic choices, health outcomes, and academic achievement ^1–3^. We operationalized self-regulation as a large heterogeneity of constituent processes that may be interrelated, including attention, set shifting, decision making, temporal discounting, response inhibition, and planning ^4,5^. As a first step in this project, we created a cognitive *ontology* of self-regulation, or an explicit specification of the entities and relationships composing a domain, which is detailed in previous work^4,5^ (data available at http://dx.doi.org/10.17605/OSF.IO/ZK6W9).

The dataset detailed in the present data descriptor was acquired to build on the cognitive ontology to measure the neural underpinnings of self regulation. We measured self regulation with a suite of converging modalities including structural MRI and functional MRI (fMRI) during tasks, rest and survey responses. The sample included 103 subjects who each completed two 90 minutes scans with nine putatively self-regulatory tasks during fMRI: attention network test ^6^ (ANT), cued task switching ^7,8^, Columbia card task ^9^, dot pattern expectancy ^10^ (DPX), delay discounting ^11^, simple stop signal ^12^, motor selective stop signal ^13^, Stroop ^14^, and a towers task ^15^. Additionally, subjects completed resting state fMRI and a set of 40 survey responses within the scanner taken from the Brief Self-Control Scale ^16^, Grit Scale ^17^, Carstensen Future Time Perspective ^18^, UPPS+P ^19^, and the Impulsiveness-Venturesomeness scale ^20^. Finally, subjects also completed anatomical scans.

This work was funded by the National Institute of Health Science of Behavior Change Commons fund (UH2DA041713). To date, this neuroimaging dataset has been used in 1 manuscript ^21^.

This data descriptor provides a description of this neuroimaging dataset, which is openly shared on OpenNeuro (https://openneuro.org/datasets/ds004636/). The shared data are organized according to the Brain Imaging Data Structure (BIDS), which is a data organization structure designed to encourage FAIR sharing and reuse of neuroimaging data ^22^.

## Methods

### Participants

Prospective participants for the study were recruited from the Stanford campus and surrounding San Francisco Bay Area using several methods including paper flyers, the Stanford Sona recruitment system, local newspapers ads, the Poldrack Lab website, online resources such as Craigslist, and through email listservs maintained by the Stanford Psychology Department. All recruited participants met the following criteria: have a minimum 8th grade education, speak English fluently, right-handed, have normal or corrected to normal vision and no color-blindness, are between 18-50 years old, have no current diabetes diagnosis, have no history of head trauma with loss of consciousness, cerebrovascular accident, seizures, neurosurgical intervention, stroke, or brain tumor, have no current major psychiatric disorders (including schizophrenia and bipolar disorder) or substance dependence, are not currently using any medication for psychiatric reasons, are not currently pregnant, and have no other contraindications to MRI.

A total of 113 participants were recruited for the study. 3 participants were dropped during their first scan session due to complications in the scanner leaving 110 participants. An additional 7 subjects did not complete their second scan session, leaving 103 subjects (64% Female, 36% Male) who completed the entire study. The sample had the following demographic distribution: 40% White, 34% Asian, 10% More than one race, 8% Black or African American, 5% Unknown, 2% Native Hawaiian or Pacific Islander, and 1% American Indian or Alaska Native.

All research activities were approved the Stanford University Institutional Review Board.

### Procedure

Scanner details and image extraction MRI data were acquired on a GE Discovery MR750 3T scanner using a Nova medical 32-channel head coil, located at the Stanford Center for Cognitive and Neurobiological Imaging (CNI).

Images were converted from DICOM data to Neuroimaging Informatics Technology Initiative (NIfTI) format using NIMSData (https://github.com/cni/nimsdata).

#### Fieldmap

A fieldmap scan was acquired with a dual-echo spiral sequence with the following parameters: TR = 700ms, TE1 = 4.545ms, TE2 = 9.1ms, flip angle = 56, 1.72 * 1.72 * 4 mm voxels. The magnitude image (magnitude.nii.gz) included in the dataset reflects the image from the first TE and the fieldmap image (fieldmap.nii.gz) shows the off-resonance frequency at each voxel in units of Hz.

#### fMRI

fMRI scans were acquired with single-echo multi-band EPI. The following parameters were used for data acquisition: TR = 680ms, multiband factor = 8, echo time = 30ms, flip angle = 53 degrees, field of view = 220 mm, 2.2 ^*^ 2.2 ^*^ 2.2 isotropic voxels with 64 slices.

#### Anatomical scans

Sagittal 3D T1-weighted scans were acquired using GE’s BRAVO sequence with TR = 7.24ms, echo time = 2.784ms, inversion time = 450ms, flip angle = 12 degrees, matrix = 256 ^*^ 256, 0.9 ^*^ 0.9 ^*^ 0.9 mm voxels. T2-weighted scans were acquired with GE’s Cube sequence with TR = 2500ms, echo time = 95.455ms, flip angle = 90 degrees, matrix = 320 ^*^ 320, 0.5 * 0.5 ^*^0.8 mm voxels.

#### Cardiac and respiratory recordings

Cardiac and respiratory data was collected with a pulse oximeter and respiration belt. We have included these raw files in the data directory within the subject and session matched with each run.

#### Participant procedure

The study consisted of 3 parts which included 2 MRI sessions and an at-home self paced online battery hosted on the Experiment Factory ^23^ (https://www.expfactory.org), which is a framework for deploying experiments in the browser. Each MRI session consisted of a 1 hour practice and setup period prior to a 1.5 hours long scanning session for a total of 2.5 hours. All MRI data was obtained at Stanford’s CNI.

Before the first MRI session, participants provided informed consent and completed a demographic survey that asked for age, sex, race, and ethnicity. This information is shared in the participants.tsv within the dataset shared.

During both scan sessions, subjects practiced the five tasks that they will do in that session in an observation room on a laptop prior to being scanned. During this practice session, a researcher read the instructions to the subject and the subject had the opportunity to ask any clarifying questions. Sessions were broken up into two groups. Group 1 included the following tasks: Stop Signal task, cued task switching task (two by two), Columbia Card Task (CCTHot), Attention Network Task (ANT), and Ward and Allport Tower task (WATT). Group 2 included the following tasks: Motor Selective Stop Signal task, Dot Pattern Expectancy task (DPX), Survey Medley, Delayed-Discounting task (DDT), and the Stroop task. Tasks were counterbalanced across subjects by employing 4 different task orderings. Separately for the two stop signal tasks, stop signal delay (SSD) was tracked during practice using a staircase algorithm and the final SSD was subsequently used as the starting SSD during the in-scanner version of the tasks. For the two stop signal tasks, response mapping for each shape were randomly decided for each participant during practice and were carried over into the in-scanner version. For the DPX task the valid cue and probe pairing was randomly chosen and used within the practice then carried over to the testing phase. For the task switching task, the color choice order and magnitude choice order were randomly chosen during practice and carried over to the testing phase. All other tasks did not counterbalance response mapping across subjects.

Scan sessions consisted of running the following sequences in order: localizer, shim, a single band reference (sbref), rest fMRI, two task fMRI runs, 2nd sbref, three task fMRI runs, and a fieldmap scan. T1-weighted scans were acquired during the first session, and T2- weighted scans were acquired during the second session. If time permitted, additional T1- weighted and T2-weighted scans were acquired during each session. All subjects were instructed to refrain from movement during scans. Task instructions were presented again during the fMRI sessions and subjects were also given instructions to respond as quickly and accurately as possible.

Participants were compensated $20/hr for their participation in the MRI sessions and $10/hr for participation in the practice section prior to scanning. Participants were also compensated $100 for their completion of a ∼10 hour online behavioral battery after the scans. This post-scan battery is briefly described below, but these post-scan data are not shared in this data release.

#### fMRI tasks

The following tasks were selected from a larger set of 36 tasks that were administered in Eisenberg et al. ^4,5^. We used a genetic algorithm to select a subset of tasks that best reconstructed the entire behavioral results across subjects in an independent dataset (see ^4^ for additional details).

All tasks made use of variable inter-trial intervals (ITIs, though see ANT) and fMRI optimized trial orderings generated through the Neurodesign python package ^24^. Neurodesign offers a four-objective optimization that allows the user to weigh four different measures of efficiency: estimation efficiency, detection power, confound efficiency, and frequency efficiency. Estimation efficiency is focused on modeling the time course of the signal, specifically using a finite impulse response function model, which was not our planned modeling approach. Detection power focuses on the power of a model using canonical hemodynamic response function-convolved regressors, which was the planned modeling approach. The other two efficiencies focus on the predictability of trials due to trial order (confound efficiency) and whether the desired trial type frequency is achieved (frequency efficiency). We chose values of 0, .1, .5 and .4 for estimation efficiency, detection power, confound efficiency and frequency efficiency, respectively, given our primary goals were to minimize predictability while retaining the desired trial frequencies with a secondary goal of maximizing detection power. Four sets of ITI’s and trial orderings were then generated by rotating both the ITI’s and trial ordering of the originally chosen Neurodesign set, each starting at a different time point within the chosen design of each task to match the four different task orders used across subjects for counterbalancing purposes. Each task used Neurodesign to power for the specific contrasts outlined below.

#### Attention network test (ANT)

The ANT ^6^ is designed to engage networks involved in three putative attentional functions: alerting, orienting, and executive control. Each trial begins with a central fixation point and a cue presented simultaneously. The cue can either be a double cue (one presented at the top and bottom of the screen, which are the two possible target locations) or a spatial cue (deterministically indicates the upcoming target location as top or bottom). A set of five arrow stimuli follows the cue at the top or bottom of the screen, and the subject must indicate via button press the direction of the arrow presented in the center of the five stimuli. The center and four flanking arrows can either point the same direction (congruent trial), or the center and the four flanking arrows can point in opposite directions (incongruent trial). Arrows can only point right or left. Responses are made with the right hand index finger if the center arrow is pointing right, and middle finger if the center arrow is pointing left.

The task consisted of 128 total trials separated into 2 blocks. Each trial consisted of a cue (100ms), fixation cross (400ms), probe (1700ms), and ITI (400ms) where a fixation is shown on the screen. The onset timing of the probe is specified in the event files included in the dataset. Responses were accepted during the probe and ITI period for each trial. Due to a coding error, this was the one task that did not include a variable inter-trial interval.

Conditions included 16 possible combinations of cue (double, spatial) * probe location (up, down) * probe direction (left, right) * condition (congruent, incongruent), each equally likely to occur. Neurodesign ^24^ was used to generate trial orders equally powering for cue, probe location, probe direction, and condition.

#### Cued task-switching task (two by two)

We used a modified cued task switching task that paired 2 different tasks with 2 different cues ^7,8^. Each trial begins with a fixation followed by a central cue instructing the subject on whether to make a magnitude or a color judgment to the subsequent probe number. The cue is either “high-low”, “magnitude”, “orange-blue”, or “color”, with the former two cues indicating one task (judge whether the subsequent probe number is greater than or less than 5), and the latter two cues indicating the other task (to judge whether the subsequent probe number is orange or blue). The cue was presented 100ms or 900ms before the target onset (with equal probabilities) and stayed on the screen throughout the probe phase. The probe was a single colored letter between 1-9 excluding 5. Responses are made with the index and middle finger of the participant’s right hand and the magnitude and color choice key mappings were randomly chosen during each subject’s practice session and used in the scanner.

The task consisted of 240 trials comprising 3 blocks of 80 trials. Each trial consisted of a fixation (500ms), cue (100, 900ms), probe (1000ms), and a blank-screen ITI (1000ms + variable ITI, see below for details). The onset timing of the probe is specified in the event files included in the dataset. Responses were accepted in the probe and ITI periods.

Conditions included cue switch (25% of trials), cue stay (25% of trials), and task switch (50% of trials). Cue switch trials are when the current trial has a different cue than the immediately preceding trial, but both indicate the same task (e.g., current trial cue is “magnitude” and previous cue was “high-low”). Cue stay trials are when the current trial has the exact same cue as the immediately preceding trial (e..g, current trial cue is “magnitude” and the immediately preceding trial cue was also “magnitude”). Task switch trials are when the current trial has a cue that indicates a different task than the task that from the immediately preceding trial (e.g., current trial cue is “magnitude” and the previous cue was “color”). Neurodesign ^24^ was used to generate both trial orders and the variable portion of the ITI sampling from an exponential distribution using a min of 0, mean of .25, and a max of 6 seconds, powering for task switch - (cue switch + cue stay) and cue switch - cue stay contrasts.

#### Columbia card task (CCTHot)

The CCTHot task ^9^ has subjects play a card game with the goal of collecting as many points by flipping over gain cards and avoiding flipping over loss cards. Each round presents a set of 6, 8, 9, 10, 12, 15, or 16 cards face down with the following information about the cards: The number of loss cards (1, 2, 3, or 5 possible), the amount they will lose if they turn over a loss card (ranging from -100 to -5), and the amount they earn with a gain card (ranging from 1 to 30). The subject can choose whether to flip over a card at random by pressing the index finger key or “cash out” and move to the next trial by pressing the middle finger key. A subject may flip over zero or as many cards as they choose, but the trial ends when they flip over a loss card. After each choice is made, the subject is shown which card they flipped over. For each round, they will be awarded the net number of points from the gain and loss cards they flipped over.

The task consists of 87 rounds split into 2 blocks of 44 and 43 rounds. Each trial is subject paced. After each round there is a feedback phase of 2250ms with an added blank-screen variable ITI (minimum of 0, mean of .2s, max of 1s). The scan was stopped after 12 minutes, so given the variable trial duration this resulted in a different number of trials per subject (*M* of completed trials across participants = 81.02, SD = 12.75, range = 32 - 87). Neurodesign was used to generate both trial orders and ITI’s ^24^.

#### Dot pattern expectancy task (DPX)

In the DPX task ^10^, subjects see a series of cue-probe stimulus pairs and make a speeded response after the probe. Each stimulus is made with a pattern of dots. There are 6 possible cues, and 6 possible probes. One cue is the “target cue” (i.e. ‘A’) and one probe is the “target probe” (i.e. ‘X’). Subjects must press one key if the target cue is followed by the target probe (i.e. ‘AX’), and another key for any other cue-probe pairing. The target cue and target probe was randomly chosen during practice and input for the test phase. Responses are made with the right hand index finger for AX pairs and middle finger for any other pair.

The task consists of 4 blocks of 40 trials for a total of 160 trials. Each trial consisted of a cue (500ms), fixation (2000ms), probe (500ms), and a blank inter-trial interval (1000ms + variable ITI). Probe responses could be made during the 500ms probe or during the inter-trial interval. The onset timing of the cue and probe are specified in the event files included in the dataset. There are four trial conditions, AX, AY, BX, and BY, with probabilities 0.55, 0.15, 0.15, 0.15, respectively. AX trials are a target cue preceding the target probe, AY trials are a target cue preceding a non-target probe, BX trials are a non-target cue preceding a target probe, and BY trials are a non-target cue preceding a non-target probe. Neurodesign ^24^ was used to generate both trial orders and the variable portion of the ITI’s. ITI were sampled from a truncated exponential distribution with a minimum of 0, a mean of .4s, and a max of 6s. We powered for AY-BY and BX-BY contrasts.

#### Delay-discounting task (DDT)

In the delayed-discounting task ^11^, subjects are presented with a dollar amount and a number of days. They are asked to choose between receiving $20 today or the larger amount of money at the presented delay in days. The amount of money ($22 - $85) and number of days (19 days - 180 days) varies across trials. Subjects were told that their performance will impact the bonus they receive. Subjects made a right hand middle finger button press for the “smaller sooner” reward of $20 and an index finger button press for the “larger later” presented reward. The task consists of 2 blocks of 60 trials for a total of 120 trials. Each Trial consists of a probe (4000ms) and a blank screen ITI (500ms + variable ITI). Responses were accepted to both the probe and the ITI.

Neurodesign ^24^ was used to generate both trial orders and ITI’s (minimum of 0, mean of .5s, max of 2.5s).

#### Stop-signal task

In the stop-signal task ^12^, subjects are presented with one of 4 “go” shapes (moon, oval, trapezoid, rectangle) and are instructed to respond to two shapes with one key press and the other two shapes with another key press. On a subset of trials, a star appears around the shape after a delay (the stop-signal delay, SSD). This indicates to the participant that they should try not to make any response on that trial. If a subject is unable to withhold their response on a stop-signal trial this is categorized as a stop failure, and if the subject makes no response then this is categorized as a stop success. During both the practice and main task, the SSD was tracked using a “1 up 1 down” staircase algorithm ^25^ in which SSD increased by 50ms after each successful stop signal trial and decreased by 50ms after each failed stop signal trial. The final SSD in practice was used as the starting SSD during the in-scanner version of the tasks.

There are a total of 125 trials, grouped into 3 blocks (41 trials in block 1 and 42 trials in each of blocks 2 and 3), with 60% of the trials being go trials and 40% being stop trials. The stop signal duration was 500ms and the initial SSD during practice was 250ms. Each trial consisted of a go stim (850ms) and an inter-trial interval that included a central fixation cross (1400ms + variable ITI). Neurodesign ^24^ was used to generate both trial orders and the variable portion of the ITI, which was sampled from an exponential distribution using a min of 0, mean of .225, and a max of 6 seconds, powering for stop - go contrast. Go responses were made with the index and middle finger of the participant’s right hand and the choice key mapping was randomly chosen during each subject’s practice session and used in the scanner.

#### Motor selective stop-signal task

The motor selective stop-signal task ^13^ is identical to the previously described stop-signal task, with the following exceptions. Instead of instructing subjects to stop whenever a stop signal occurs, subjects are instructed to only stop if a stop signal occurs and they were going to make one of their two responses (the “critical” response), but not if a stop signal occurs and they were going to make the other response (the “non-critical” response). SSD was adjusted only on critical stop trials, and the SSD on noncritical stop trials was yoked to the SSD on the critical stop trials. Responses are made with the index and middle finger of the participant’s right hand and the choice key mapping is randomly chosen during each subject’s practice session and used in the scanner. Critical stop trial stimuli are also randomly chosen during the subject’s practice session and used in the scanner.

The task consists of 5 blocks of 50 trials for a total of 250 trials. 60% of trials were go trials, 20% critical stop trials, and 20% noncritical stop trials. We powered for the following contrasts: Noncritical stop - noncritical no-signal and critical stop - noncritical stop.

#### Stroop task

In the Stroop task ^14^, subjects are presented with a color word (e.g., “red”) written in ink that either matches the word (congruent, e.g., “red” in red) or does not (incongruent, e.g., “red” in blue). Subjects are instructed to quickly and accurately respond via keypress to the ink color of the word is. The color of the word and the written word could both be red, blue or green. The participants were asked to respond with index, middle, and ring finger for each of these colors.

The task consists of 2 blocks of 48 trials for a total of 96 trials. Each trial consists of a probe (1500ms) and an inter-trial interval that includes a central fixation cross (500ms + variable ITI). There were 48 congruent and 48 incongruent trials. Neurodesign ^24^ was used to generate both trial orders and the variable portion of the ITI sampling from an exponential distribution using a min of 0, mean of .2, and a max of 6 seconds, powering for incongruent - congruent.

#### Ward and Allport tower task (WATT)

In the WATT ^15^ subjects are presented with stacked balls in a specific initial configuration on three pegs, and they are asked to move the stacked balls to a different target configuration using the least amount of moves.

The task consists of 3 blocks of 16 rounds for a total of 48 rounds. These moves are not timed and the round ends when the subject moves the ball to match the target configuration shown on the screen. All trials were designed to be completed in 3 steps, where a step is defined by taking a ball off the board then placing it on a different peg. After the participant correctly solves each round there is a feedback phase that consists of feedback shown for 1000ms and a blank screen (1000ms + variable ITI). Some subjects did not complete the full 48 rounds, as there was a max scan time of 10 minutes and the task was ended after 10 minutes even if all trials were not completed (*M* of completed rounds across participants = 45.35, SD = 4.99, range = 28 - 48). Participants practiced with trials in which the target configuration had all three balls positioned on the same peg (Unambiguous). The main test phase consisted of trials where the target configuration had one ball on one peg and two balls on another peg (Partially Ambiguous). Trials included conditions in which a ball had to be moved out of the way to reach the target position (with intermediate step) and in which all balls could directly be moved to the target position (without intermediate step). Both practice and test phase were scanned.

Neurodesign ^24^ was used to generate both trial orders and the variable portion of the ITI sampling from an exponential distribution using a min of 0, mean of .9, and a max of 6 seconds, powering for conditions shown above.

#### Survey medley

In addition to the above 9 traditional cognitive tasks, subjects also completed 40 survey questions that were answered within the scanner. We scanned the following three complete surveys: the Grit Scale ^17^ (8 questions), Brief Self-Control Scale ^16^(13 questions), and the Carstensen Future Time Perspective^18^ (10 questions, Carstensen). We also selected a subset of items from the Impulsiveness-Venturesomeness^20^ (3 questions) and UPPS+P impulsivity^19^ (6 questions) questionnaires.

Each trial presented the survey question for 8500ms and was followed by a blank screen for 500ms plus a variable ITI (minimum of 0, mean of 1s, max of 5s). Neurodesign (Durnez et al., 2018) was used to generate both trial orders and ITI’s.

#### Post-scan battery

Upon completion of the scanning sessions participants were also given an at-home behavioral battery to do on their own time that consists of 63 tasks and surveys, matching the full battery from Eisenberg et al. ^5^. These data are not included within this data release and are not described in detail in this data descriptor.

### Data Records

The dataset is hosted on OpenNeuro under accession number ds004636. The files are organized in the Brain Imaging Data Structure (BIDS) format, including sidecar JSON files with detailed explanation of MRI scan protocol parameters as well as task parameters. Functional scans are labeled as follows:

- ANT: Attention network test
- CCTHot: Columbia card task
- DPX: Dot pattern expectancy
- WATT3: towers task
- discountFix: delayed discounting
- motorSelectiveStop: Motor selective stop-signal task
- rest: Resting state (eyes open)
- stopSignal: Stop-signal task
- Stroop: Stroop task
- surveyMedley: 40 survey questions
- twoByTwo: Cued task switching task

The participants.tsv file contains age and sex of each participant, and quality_control.csv contains notes on suggested exclusions based on behavior and neuroimaging data quality assurance (described below).

Cardiac and respiratory recordings are included as gzip compressed .tsv files for each functional run. The cardiac data files are named in the following way: sub-{subject_number}_ses- {session_number}_task-{task_name}_run-{run_number}_recording-cardiac_physio.tsv.gz. The respiratory data files are named in the following way: sub-{subject_number}_ses- {session_number}_task-{task_name}_run-{run_number}_recording-respiratory_physio.tsv.gz. A JSON file with relevant metadata is included in the main directory. As noted in the JSON file, cardiac and respiratory data were recorded starting from 30 seconds before the functional time series started.

### Derivatives

All anatomical and functional scan images were run through MRIQC version 22.0.6 ^26^ (RRID:SCR_022942). T1-weighted and T2-weighted images were defaced using pydeface version 2.0.0 ^27^ prior to being put through MRIQC. MRIQC reports are shared as Hypertext Markup Language (html) files and image quality metrics (IQM) metadata are shared as JSON files in the derivatives/mriqc directory within the root BIDS directory. BIDS derivatives are included as a directory on OpenNeuro ds004636.

### Behavioral data

Raw task behavior during scans were collected using Experiment Factory ^23^(https://www.expfactory.org) based on jsPsych 5 ^28^. These data include the in-scanner practice portion of the task that was not scanned, if it was included in task structure. Raw data were cleaned and processed to create event files using python scripts (https://doi.org/10.5281/zenodo.8350250). All raw and cleaned behavioral data are shared in an OSF repository (https://doi.org/10.17605/OSF.IO/SJVNW). Data are organized by subject number and task name.

### Technical Validation

All collected data are included in the current data release. We have completed quality assurance of the data that is described in detail in the immediately following sections, and we recommend that certain subjects and scans be excluded, but we recognize that exclusion decisions can depend on the goals of each project, so we have decided to include all available data in the data release so that data users can make exclusion decisions that align with their use cases.

### Behavioral Quality Control

We completed the following behavioral quality control, and we have provided annotations of these quality control recommendations in suggested_exclusions.csv on OpenNeuro.

Falling outside the normal range on any of the following criteria was sufficient for that subject’s data to be recommended for exclusion. In the stop-signal tasks, stop success rate being below .25 or above .75, in line with consensus recommendations ^29^. In all tasks, having an omission rate of over .5. Subjects missing more than half the tasks had their entire dataset (all tasks) recommended for exclusion. We ran a binomial test on noncritical signal trials in motor selective stopping to ensure that subjects were making responses on more than half of trials. If they failed this binomial test, they were recommended for exclusion as it indicated they were treating the task as a simple stop signal task. Last, we looked to see if any subjects omitted a majority of their responses at the beginning or end of any scans, but this did not identify any additional subjects.

We also evaluated descriptive statistics for each subject on each task, including their accuracy, RT, and other key dependent variables (e.g., range of SSDs for the stop-signal task), for aberrant behavior. This resulted in the recommended exclusion of 11 tasks across 6 subjects, mostly for accuracy that was at or below chance in at least one condition. Details of each of these 11 decisions can be found in the subjective_rating_descriptions.json within the BIDS directory.

As a result of this quality assurance effort, we recommend that 12 subjects be completely excluded because performance in at least 5 of their 9 speeded tasks (i.e., all but the survey medley) were deemed unsatisfactory. Additionally, we identified 22 subjects who had satisfactory behavior on at least 5 of their 9 tasks but we recommend excluding their data for 1 to 4 of their tasks based upon performance. Therefore, of the 103 dataset, 68 are judged to be both complete and have satisfactory behavioral data on all tasks.

The sample after our suggested behavioral exclusions (63% Female, 37% Male; Age 19- 44, mean: 25) consists of the following demographic distribution: 45% White, 31% Asian, 9% More than one race, 9% Black or African American, 5% Unknown, and 1% Native Hawaiian or Pacific Islander.

### Neuroimaging Analyses and Quality Control

Imaging files were converted from DICOM data to Neuroimaging Informatics Technology Initiative (NIfTI) format using NIMSData (https://github.com/cni/nimsdata). T1-weighted anatomical images and BOLD contrast (fMRI) images were processed with MRIQC ^26^ (RRID:SCR_022942) to extract the various image quality metrics listed below. MRIQC image quality metrics outputs are included in OpenNeuro derivatives. Scripts used to generate the attached figures are included in the following repository: 10.5281/zenodo.8350250.

### Anatomical Scans

Anatomical T1w images were defaced using pydeface version 2.0.0 (Gulban et al., 2019) and the defaced images were used to calculate the following measures for quality control using MRIQC ^26^ (RRID:SCR_022942). See Figure 1 for T1w quality assurance metrics and Table 1 for an explanation of the quality control metrics.

**Table 1.** Description of QC measures of T1-weighted anatomical data.

| Image Quality Metric (Label) | Description |
| --- | --- |
| Artifact detection ($qi\_1$) | Detects artifacts from motion, blurring or ghosting<sup>30</sup>. Lower values are better. |
| Contrast-to-noise ratio (cnr) | Measure of contrast between white and gray matter<sup>31</sup>. |
| Entropy focus criterion (efc) | Proxy for ghosting or blurriness caused by head motion<sup>32</sup>. Lower values are better. |
| Foreground-background energy ratio (fber) | Average value of energy within the head compared to regions outside of the head<sup>33</sup>. Higher values are better. |
| Gray matter signal (summary_gm_mean) | Average signal of gray matter. |
| Signal-to-noise ratio (snr_total) | Relative measure of background noise. Higher values are better. |

**Table 2.** Description of QC measures of BOLD data.

| Image Quality Metric (Label) | Description |
| --- | --- |
| Ghost to Signal Ratio (gsr_y) | Measure of signal intensity in areas along encoding direction y compared to signal intensity inside the brain<sup>34</sup>. Lower values are better. |
| DVARS (dvars_vstd) | Measure of signal changes from volume to volume. Lower values are better. |
| Framewise displacement (fd_mean) | Measure of motion throughout the scan measured in millimeters. Lower values are better. |
| Foreground-background energy ratio (fber) | Average value of energy within the head compared to regions outside of the head<sup>33</sup>. Higher values are better. |
| Temporal SNR (tsnr) | Relative measure of background noise across time course. Higher values are better. |
| Signal-to-noise ratio (snr) | Relative measure of background noise. Higher values are better. |

**Figure 1.**
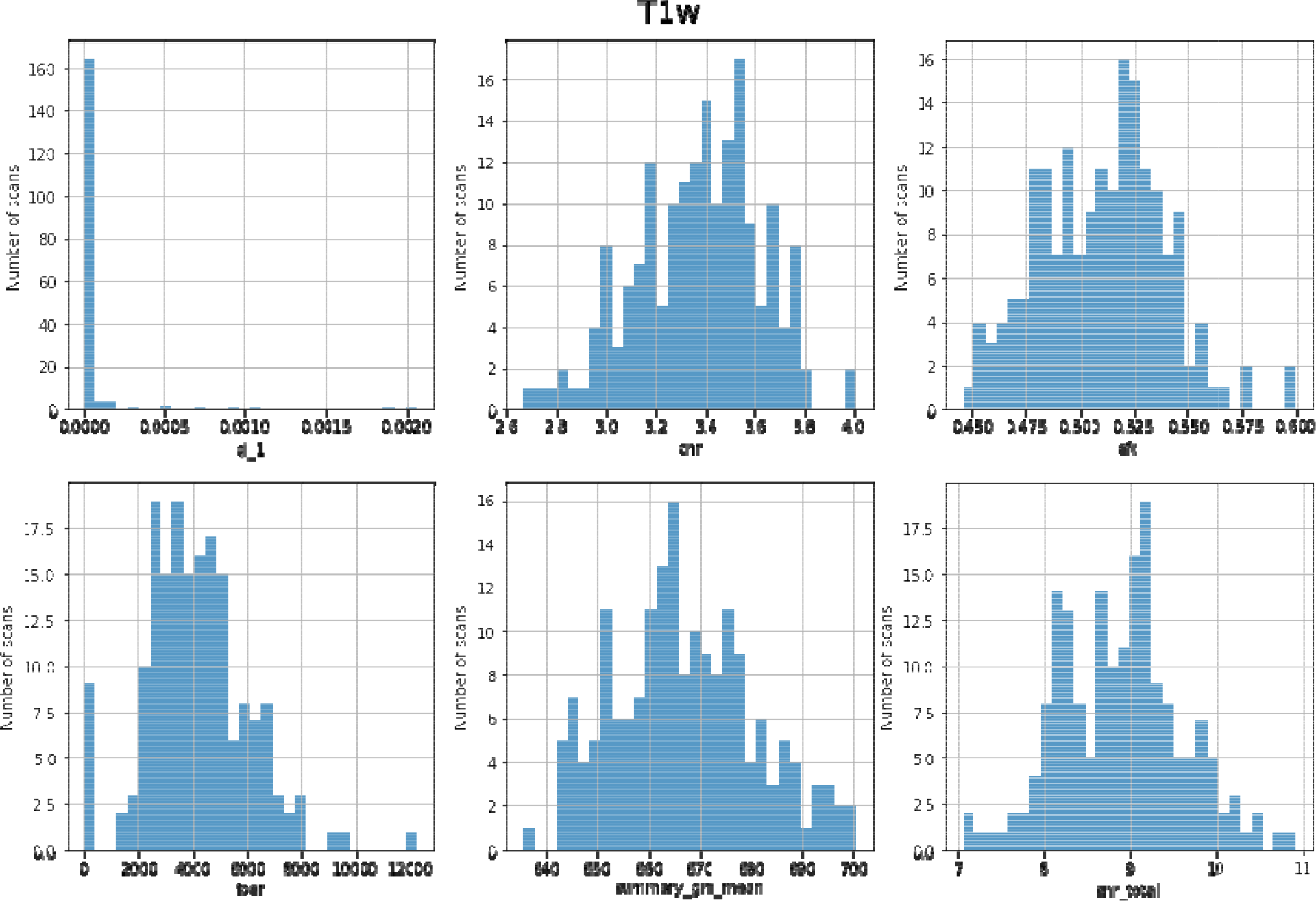
Distribution of QC measures of T1-weighted data.

### Functional BOLD scans

The following quality control metrics were computed using MRIQC for each fMRI scan.

MRIQC reports were visually inspected for visible artifacts such as aliasing, distortion, and ghosting. Specific scans with MRIQC reports that did not pass inspection are noted in quality_control.csv within the dataset. See Figure 2 for distributions of quality control measures across all functional scans (task and rest) and Table 1 for an explanation of the quality control metrics. Quality control measures were highly similar across functional scans, so we include all task and rest data aggregated in Figure 2 (gsr_y (*M* = 0.038, SD = 0.001, range = 0.037 - 0.040), dvars_vstd (*M* = 0.986, SD = 0.012, range = 0.967 - 1.010), fd_mean (*M* = 0.108, SD = 0.008, range = 0.092 - 0.120), fber (*M* = 11672.360, SD = 379.258, range = 11149.039 - 12441.484), tsnr (*M* = 36.842, SD = 2.646, range = 33.108 - 42.351), snr (*M* = 3.281, SD = 0.020, range = 3.247 - 3.310)).

**Figure 2.**
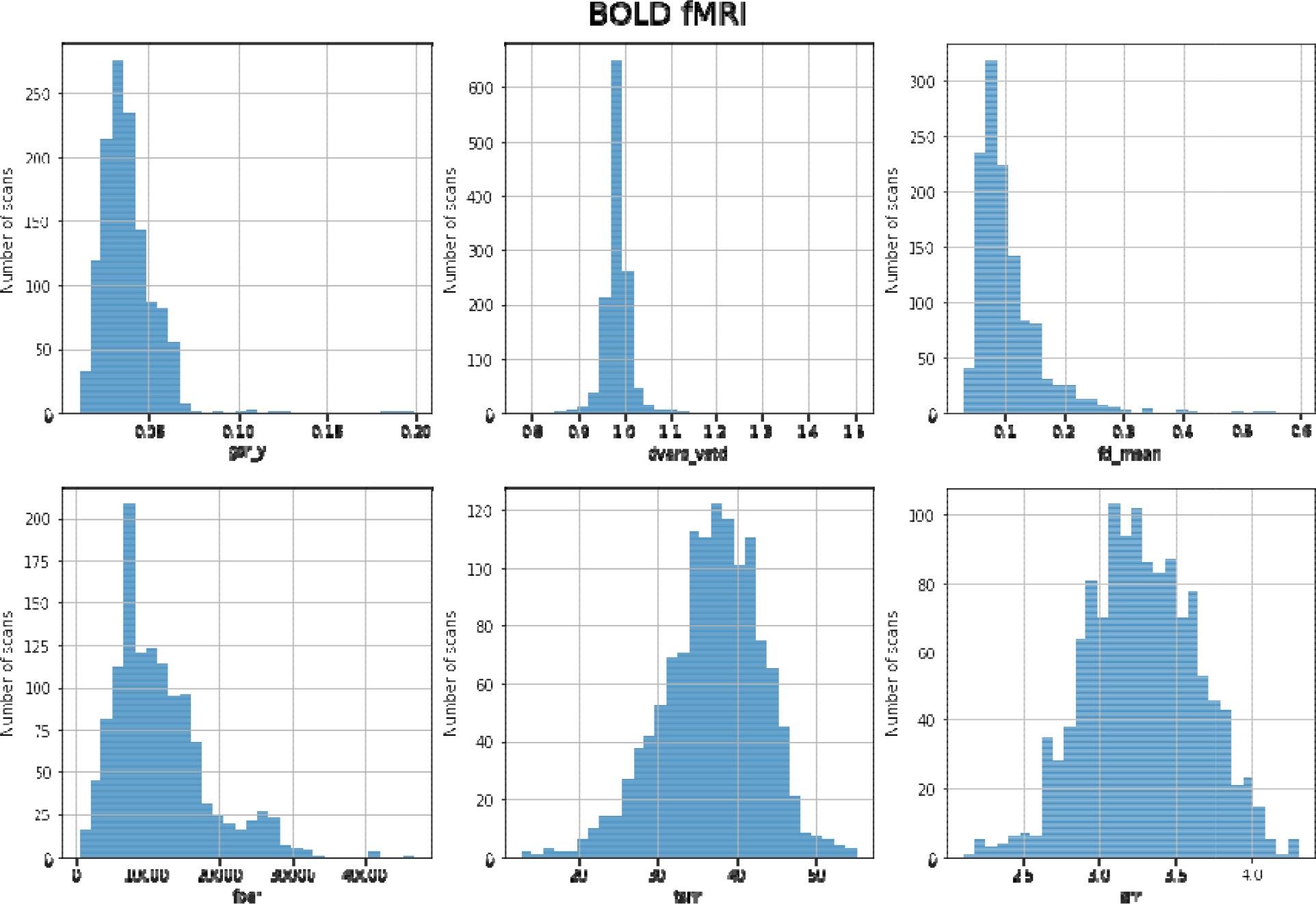
Distribution of QC measures for BOLD functional scans.

### Usage Notes

All data are made available under a Creative Commons CC0 public domain dedication, which places no restrictions on the reuse or redistribution of this dataset. Users of the data should follow the ODC Attribution-Sharealike Community Norms (https://opendatacommons.org/norms/odc-by-sa/), by sharing any derivative works with an equally open license, and by crediting the creators of this dataset by citing this data descriptor. All code released in relation to this publication is released under an MIT License.

## Code Availability

Neuroimaging data is available on OpenNeuro under accession number ds004636 (https://openneuro.org/datasets/ds004636/). Raw behavioral data of fMRI tasks is available on OpenScience Framework (https://doi.org/10.17605/OSF.IO/SJVNW). All scripts used to create event files and to create figures in this manuscript are available in GitHub repository Self_Regulation_Ontology_Neuro (https://doi.org/10.5281/zenodo.8350250).

## Acknowledgements

This work was supported by the National Institutes of Health (NIH) Science of Behavior Change Common Fund Program through an award administered by the National Institute for Drug Abuse (NIDA) (UH2DA041713; PIs: Marsch, LA & Poldrack, RA).

## Author contribution statement

Eisenberg, I. W., Enkavi, A. Z., Li, J. K., and Bissett, P. G., acquired the data. Bissett, P. G., Eisenberg, I. W., Shim, S., Rios, J. A. H., Jones, H. M., Hagen, M. P., Enkavi, A. Z., Li, J. K., Mumford, J. A., & Poldrack, R. A. quality assured and analyzed the data. Poldrack, R. A., Marsch, L. A., & MacKinnon, D. P., acquired funding for the research and guided the data acquisition.

